# CNS-Tau Specific Antibodies Illuminate Disease Signatures Across Tauopathies

**DOI:** 10.1101/2025.09.14.676119

**Authors:** Lei Liu, Yin Zheng, Luke Slominski, Qimin Quan, Adriana Saba, Jean-Pierre Bellier, Han-Bin Yoo, Andrea Roman, Elizabeth L. Hennessey, Samadhi P. Wijethunga, Elif L. Okyay, Michael B. Miller, Hyun-Sik Yang, Dennis J. Selkoe, Jasmeer P. Chhatwal

## Abstract

**Background:** Alternative splicing of the MAPT gene produces distinct tau isoforms in the central and peripheral nervous systems (CNS and PNS), yet their respective biological and pathological roles remain poorly understood. Recent studies suggest that CNS-tau may play a key role in amyloid-β associated neurodegeneration in Alzheimer’s disease (AD), but the absence of isoform-specific tools has limited both mechanistic insight and biomarker development. We aimed to develop and validate CNS-tau–specific monoclonal antibodies and assess their utility in neuropathology and fluid-based biomarker assays across AD and primary tauopathies.

**Methods:** We generated six recombinant rabbit monoclonal antibodies targeting a CNS-tau–specific sequence encoded by exons 4 and 5. Specificity and affinity were evaluated via biolayer interferometry, immunoblotting, and tau-expressing HEK293 models. The lead clone, LL-T-1-1, was tested in postmortem brain sections from AD (n = 23), progressive supranuclear palsy (PSP, n = 3), and corticobasal degeneration (CBD, n = 3). A second clone, LL-T-1-5, was optimized for use in plasma assays via an ultrasensitive nanoneedle platform, LoD < 1 pg/ml.

**Results:** LL-T-1-1 showed nanomolar affinity for CNS-tau and no cross-reactivity with PNS-tau. It selectively labeled dystrophic neurites in AD and all hallmark tau lesions in PSP and CBD without antigen retrieval. LL-T-1-5–based plasma assays revealed CNS-tau levels significantly correlated with cognitive scores (MMSE and QDRS) and differentiated impaired from unimpaired individuals.

**Conclusions:** CNS-tau–specific antibodies LL-T-1-1 and LL-T-1-5 provide new tools for neuropathology and fluid biomarker development across tauopathies.

## Introduction

The rapid progress of plasma biomarkers for Alzheimer’s disease (AD) since 2020 has paralleled the clinical advancement of disease-modifying therapies (DMTs) such as lecanemab and donanemab. A new generation of biomarkers—including oligomeric Aβ^1^, CNS-specific tau isoforms^2,3^, and various phosphorylated tau species^4^—now holds promise for improving diagnosis, staging, prognosis, and therapeutic monitoring. However, the development of these biomarkers has largely been empirical, with limited consideration of the underlying molecular and pathophysiological frameworks that define their origins and significance.

To help address this gap, we developed a suite of highly specific recombinant antibodies targeting distinct tau proteoforms, including multiple phosphorylation sites associated with the proline-directed kinase motif (pTPP)^5^. Our prior work demonstrated that sites such as pT175, pT181, pT217, and pT231 exhibit strong intercorrelation in AD brains, emphasizing that the overall phosphorylation burden—rather than individual phospho-epitopes—may more accurately reflect tauopathy severity and progression^5^.

In the present study, we extend this approach by focusing on isoform-specific tau biology, particularly the distinction between central nervous system (CNS) and peripheral nervous system (PNS) tau splicing variants. Despite early recognition of distinct CNS and PNS tau isoforms^6^, there has been a lack of isoform-specific detection tools. Recent studies have implicated CNS-tau–specific signals in amyloid-β-associated neurodegeneration^3^, but no commercial antibodies with proven isoform-level specificity in fluid or tissue contexts. To fill this unmet need, we developed a panel of recombinant rabbit monoclonal antibodies targeting CNS-tau–specific sequences encoded by the junction of exons 4 and 5.

In summary, this study introduces and validates the first CNS-tau–specific antibody suitable for use in both fluid-based and tissue-based assays. Our findings suggest that CNS-tau isoform– specific signals reflect disease-relevant neuropathology and hold promise for advancing biomarker development across multiple human tauopathies.

## Material and Methods

### Participants

Participants were enrolled from the Davis Memory and Aging Cohort (MAC), a longitudinal observational study at Mass General Brigham that recruits individuals aged 45 years and older across a spectrum of cognitive functioning. For this analysis, we included 72 participants with complete data on self-reported cognitive decline, cognitive testing, and study partner–reported functional assessments. The mean age was 68.7 years (SD = 9.8). Cognitive status was categorized using validated Quick Dementia Rating System (QDRS) thresholds. All participants completed a brief three-item self-report questionnaire adapted from the NIA-AA research framework, underwent the Mini-Mental State Examination (MMSE), and had an accompanying study partner complete the QDRS.

### Generation of human Tau-441 and Tau-758 expression vectors

The coding sequences of wild type human Tau-441 and Tau-758 isoforms were synthesized by IDT and subcloned into a pcDNA3.1 vector. Vectors were sequenced from 5’ and 3’ ends for validation.

### Tissue culture and transfection of adherent cells

Adherent HEK cells were cultured in complete growth medium: Dulbecco’s Modified Eagle’s Medium (DMEM) supplemented with 10% fetal bovine serum (FBS), two mM L-glutamine, 10 Units/mL penicillin and 10 μg /mL streptomycin. Adherent HEK cells were seeded in 24-well dishes at a density of 5 x 10^5^ cells per well for transfection. Transfection was carried out with PEI-max reagent (Polysciences). Cells were cultured for 48 hr before the lysates were harvested for immunoassay and western blot.

### Antibodies

BT-2 (RRID: AB_10975238) was from Thermofisher. Tau-12 (RRID: AB_2715842), 77G7 (RRID: AB_2564801) and Tau-13 (RRID: AB_291452) were from Biolegend. D1M9X (RRID: AB_2783844) was from Cell Signaling Technology. CNS specific rabbit monoclonal antibodies were generated using a custom-designed synthetic peptide antigen conjugated to KLH at the N terminus (LifeTein). After 8 boosters of immunization, peripheral blood mononuclear cells (PBMCs) are isolated from immunized rabbits and affinity enriched with antigen used for immunization through Fluorescence-activated Cell Sorting (FACS) into single B cells. Single B cells were used for sequencing of variable regions of antibodies, and the sequences were sub-cloned into expression vector for antibody production. From 99 sequence-derived antibodies, we selected 6 best candidates for follow-up analysis, which became the LL-T-1 series.

### Recombinant tau protein production

DNAs coding tau protein 1-210 (Tau 441 isoform) and 1-500 (Tau 758 isoform) were sub-cloned in pTXB1 plasmid (New England Bio, USA) and transformed into *Escherichia coli* BL21(DE3) strain for expression. Transformed cells were grown at 37°C in 500 ml LB medium with 100 μg/ml ampicillin at 300 rpm. When A600 reached 0.6–1.0, IPTG was added to a final concentration of 0.4 mM to induce the expression. After further incubation at 37°C for 2 h, the cells were collected by centrifugation at 3000 g for 15 min at 4°C. The cell pellet was resuspended in 10 ml of lysis buffer (20 mM histidine, pH 6.0, 50 mM NaCl, 1 mM EDTA, 5 mM DTT, 0.1 mM PMSF, and 5% (v/v) glycerol), quickly frozen in liquid nitrogen, and stored at 80°C until use. Large-scale purification of recombinant tau proteins was performed at 4°C on a fast protein liquid chromatography (FPLC) system (ÄKTA, Cytiva) using a combination of anion-exchange chromatography (AIEXC) and size exclusion chromatography (SEC).

### Biolayer Interferometry (BLI)

Rabbit antibodies were immobilized onto protein-A biosensor (Sartorius, 18-5010) to density ∼1.0 nM using an Octet RED96e (ForteBio). After equilibrium, binding to 100 nM CNS-tau and PNS-tau recombinant proteins, buffered in pH 7.4 PBS buffer with 0.1% BSA, was performed using 180 s association and 180 s dissociation at 25°C. Binding affinity was fitted using a 1:1 binding model.

### Electrophoresis and WB

Samples were loaded onto 4-12% Bis-Tris gels (SurePAGE, Genscript) using MES-SDS running buffer, transferred to nitrocellulose membranes, and probed for various proteins using standard WB. For visualization, membranes were treated with HRP-labeled secondary antibodies (JCI) and then with DAB substrate (TCI) with intensification by nickel ammonium sulfate. Blot images were scanned with a well-calibrated Perfection V850 Pro Photo Scanner (Epson).

### Tau Immunoassay (MSD)

HEK cell lysates were diluted with 1% BSA in wash buffer (Tris-buffered saline [TBS] supplemented with 0.03% Tween). For all assays, each well of an uncoated 96-well multi-array plate (MesoScale Discovery) was coated with 30 μL of a PBS solution containing 1 μg/mL of Tau-13 capture antibody to the N-terminus region of Tau (epitope aa 15-25) and incubated at RT overnight. The aforementioned detection antibody solutions were prepared to have biotinylated monoclonal antibodies (LL-T-1 series and certain commercial antibodies), plus 100 ng/mL Streptavidin Sulfo-TAG (MesoScale Discovery, #R32AD-5) and 1% BSA diluted in wash buffer. Following overnight incubation at RT, 50 μL/well of the sample, followed by 25 μL/well of detection antibody solution were incubated for 2 hr at RT with shaking at 850 rpm, and washing of wells with wash buffer between incubations. The plate was read and analyzed according to the MSD manufacturer’s (Meso Scale Diagnosis) protocol.

### Immunostaining of human brain sections

The Harvard Brain and Tissue Resource Center (HBTRC, at McLean Hospital) and the BWH Neuropathology tissue repository provided autopsied brain sections. Formalin-fixed, paraffin-embedded sections (5 μm) of the dorsolateral prefrontal cortex BA9 (DLPFC) from 23 subjects covering the entire span of AD neuropathology from Braak stages 0 to 6 and 6 subjects having non-AD tauopathy were used in this study. The paraffin sections were rehydrated, and endogenous peroxidases were inactivated by treatment with 0.3% H_2_O_2_ in PBS containing 0.03% Triton X-100. The sections were incubated with primary antibodies at 4°C overnight and then with biotinylated secondary antibodies for 1 hr. For visualization, sections were treated with avidin/biotin-HRP complex (Vector) and then with DAB substrate (TCI) with intensification by nickel ammonium sulfate. Photomicrographs were taken with an CX33 microscope (Olympus) equipped with Mlchrome 5 Pro camera (Tucsen), and brightness/contrast/threshold were adjusted with ImageJ 1.54p (NIH).

### CNS-Tau plasma assay (Nanoneedle technology)

LL-T-1 series antibodies were paired with Tau-12 on the Nanoneedle assay. 5 μg/ml LL-T-1 series capture antibody was incubated on the nanoneedle plate overnight, followed by blocking buffer for 1 hr. 10 μL plasma samples were thawed and diluted 3x into the dilution buffer and measured in triplicate on the nanoneedle plate. The sample incubation was 2 hr, followed by wash steps and incubation with biotinylated Tau-12 antibodies for 1 hr. A mass amplifier solution (NanoMosaic) was then incubated for 30 min and induced stronger color change on the nanoneedles. The nanoneedle plate was imaged using the Tessie ^™ □^ instrument (NanoMosaic).

## Results

### Generation of highly specific antibodies against CNS-Tau isoforms

To investigate the biological and pathological significance of CNS-specific tau isoforms, we developed a panel of antibodies with high specificity for CNS-tau and no cross-reactivity with PNS-tau. Although a previous study reported a home-brew sheep monoclonal antibody raised via hybridoma screening^2^, the only commercially available reagent—offered by Roboscreen GmbH—lacked rigorous analytical validation. Given the distinct B-cell ontogeny and elongated CDR3 of rabbits^7^, we selected a rabbit monoclonal antibody platform for its superior affinity and epitope recognition. We immunized rabbits with a synthetic peptide corresponding to residues 110–130 of the 2N4R tau isoform (Tau 441), which spans exons 4 and 5 of the *MAPT* gene—a junctional sequence uniquely present in CNS tau but absent from PNS tau (Figure 1A). Through a single B-cell isolation and recombinant expression workflow, we generated six monoclonal antibodies (LL-T-1 series).

**Figure 1.**
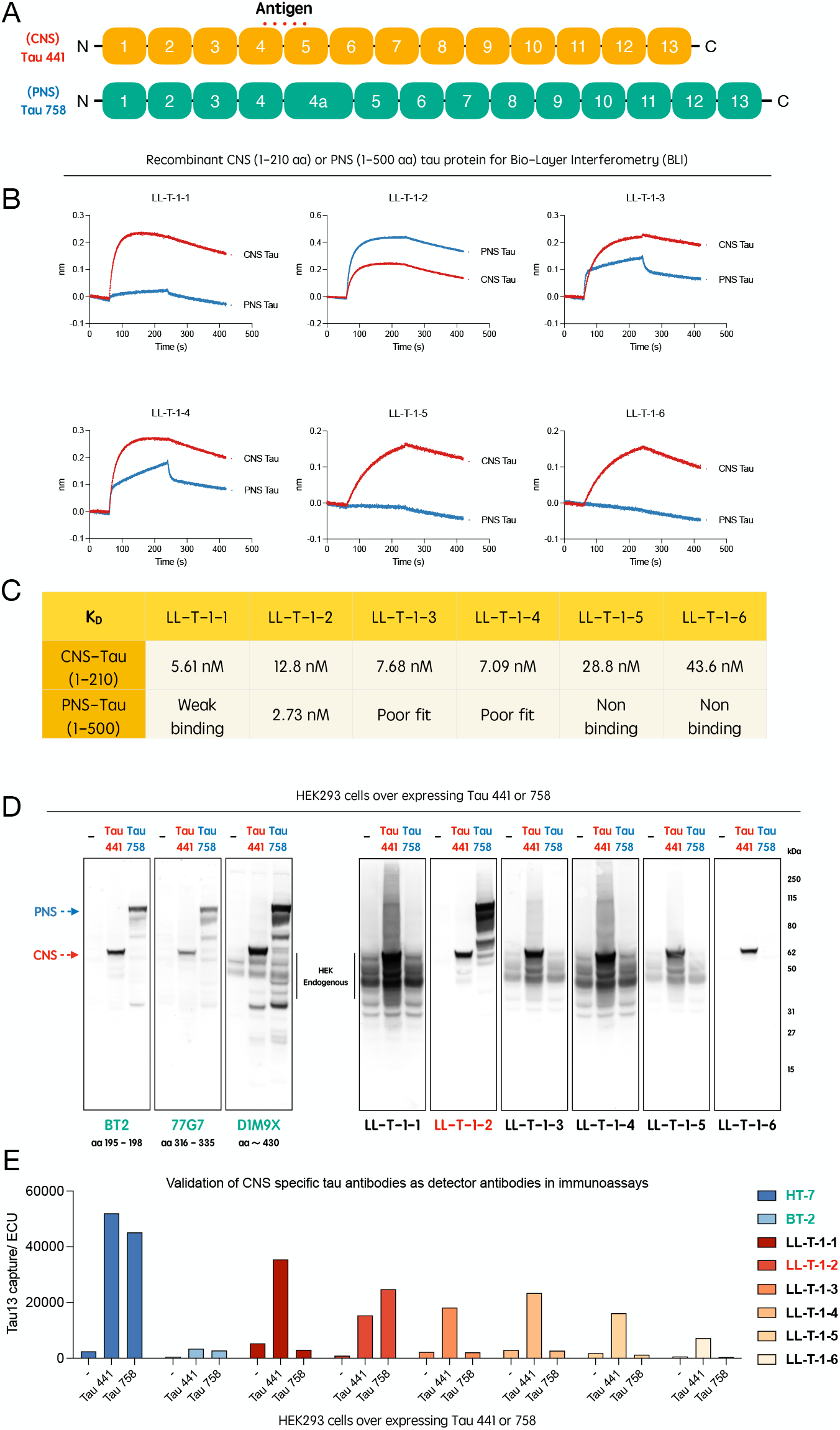
Generation and validation of CNS-tau–specific antibody series LL-T-1. (A) Schematic representation of the tau protein, highlighting the epitope region unique to CNS-tau isoforms (junction sequence between protein encoded by exons 4 and 5). (B) Sensorgrams from biolayer interferometry (BLI) showing binding kinetics of LL-T-1 antibodies to recombinant CNS- and PNS-tau proteins. (C) Calculated equilibrium dissociation constants (Kd) calculated from BLI data for all six antibodies against CNS- and PNS-tau. (D) Immunoblots of lysates from HEK293 cells transfected with full-length CNS-tau (Tau-441) or PNS-tau (Tau-758), probed with commercial tau antibodies or LL-T-1 clones. (E) Isoform-selective immunoassays using Tau-13 as the capture antibody and commercial antibodies or LL-T-1 clones as detectors.

Binding specificity and affinity were assessed by biolayer interferometry (BLI) using recombinant CNS-tau and PNS-tau. Among the six clones, LL-T-1, LL-T-5, and LL-T-6 showed strong preference for CNS-tau, while LL-T-2 favored PNS-tau (Figure 1B). LL-T-1-1 demonstrated nanomolar affinity for CNS-tau with minimal cross-reactivity (most left panel of Fig. 1C), making it the lead candidate. In immunoblots using HEK293 cells transfected with either CNS-tau (Tau-441) or PNS-tau (Tau-758), five out of six clones showed strong CNS-tau specificity, in contrast to the non-selective commercial antibodies BT2, 77G7, and D1M9X (Figure 1D). Notably, LL-T-1-1, -3, -4, and -5 detected endogenous tau at concentrations below 100 ng/mL. These findings were consistent with the BLI-derived affinity rankings. We further validated antibody performance in a sandwich immunoassay format, using Tau-13 as the capture antibody and the LL-T-1 clones as detectors. Commercial antibodies such as HT7 and BT2 detected both isoforms equally, while five out of six LL-T-1 clones selectively detected CNS-tau (Figure 1E).

### LL-T-1-1 reveals disease-relevant Tau pathology across tauopathies and Braak stages

Following extensive characterization, we applied LL-T-1-1 to paraffin-embedded human brain sections from BA9 across three tauopathy cohorts: sporadic Alzheimer’s disease (AD, n = 23), progressive supranuclear palsy (PSP, n = 3), and corticobasal degeneration (CBD, n = 3). Remarkably, LL-T-1-1-stained pathological tau structures at low concentrations (100 ng/mL) without antigen retrieval (Figure 2). In AD brains, LL-T-1-1 preferentially labeled dystrophic neurites (Figure 2A.1), and to a lesser extent, neurofibrillary tangles (Figure 2A.2) and corpora amylacea (Figure 2A.3). This staining pattern contrasts with phospho-tau antibodies like AT8 or pT217, which equally label tangles and dystrophic neurites^5^. As shown in Figure 2B, double immunofluorescence with AT8 and LL-T-1-1 revealed perfect co-labeling of dystrophic neurites, but not all tangles—some of which remained unlabeled by LL-T-1-1, as highlighted within the white dotted circle in the lower panel. In PSP and CBD, LL-T-1-1 identified hallmark tau lesions^8^ including globose tangles (Figure 2C.4), tufted astrocytes (Figure 2C.5), oligodendroglial coiled bodies (Figure 2C.6), ballooned neurons (Figure 2D.7), astrocytic plaques (Figure 2D.8), and neuropil threads (Figure 2D.9). This broad reactivity suggests that the antibody recognizes a shared epitope across tau conformers in both amyloid-β-associated and primary tauopathies. To determine the relationship between LL-T-1-1 staining and disease progression, we evaluated its signal across Braak stages 0–6 within the same AD cohort (n = 23). In lower stages (0–3), staining was weak, likely reflecting its lack of recognition of physiological CNS-tau expression. By Braak stages 4–6, strong labeling appeared in neuronal threads, dystrophic neurites, and neuritic plaques (Figure 3B). Densitometric analysis confirmed a Braak stage–dependent increase in CNS-tau–positive area (Figure 3A), demonstrating the antibody’s sensitivity to AD disease stage.

**Figure 2.**
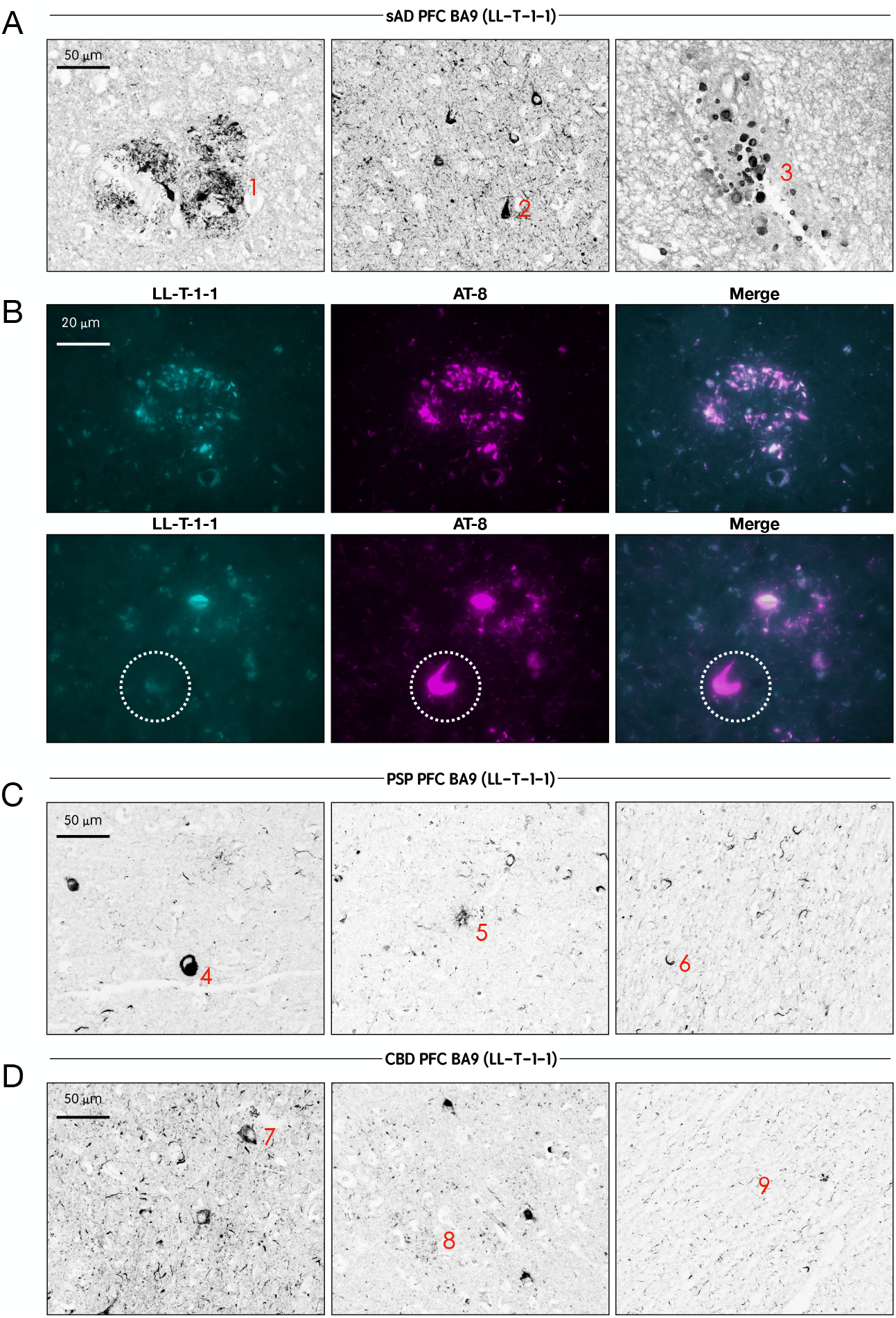
LL-T-1-1 preferentially labels dystrophic neurites in AD and robustly detects classic tau lesions in PSP and CBD brains. (A) High-magnification images of LL-T-1-1 immunostaining in Alzheimer’s disease (AD) brain sections reveal abundant labeling of dystrophic neurites (1), neurofibrillary tangles (2), and corpora amylacea (3) (red labels). (B) High-magnification immunofluorescence images showing LL-T-1-1 (cyan) and AT8 (purple) co-staining. LL-T-1-1 and AT8 co-label dystrophic neurites, while AT8 additionally stains neurofibrillary tangles that are not consistently recognized by LL-T-1-1, as highlighted by the white circle. (C) LL-T-1-1 immunostaining in progressive supranuclear palsy (PSP) brain sections demonstrate robust labeling of globose tangles (4), tufted astrocytes (5), and oligodendroglial coiled bodies (6) (red labels). (D) In corticobasal degeneration (CBD) brain sections, LL-T-1-1 highlights ballooned neurons (7), astrocytic plaques (8), and neuropil threads (9) (red labels).

**Figure 3.**
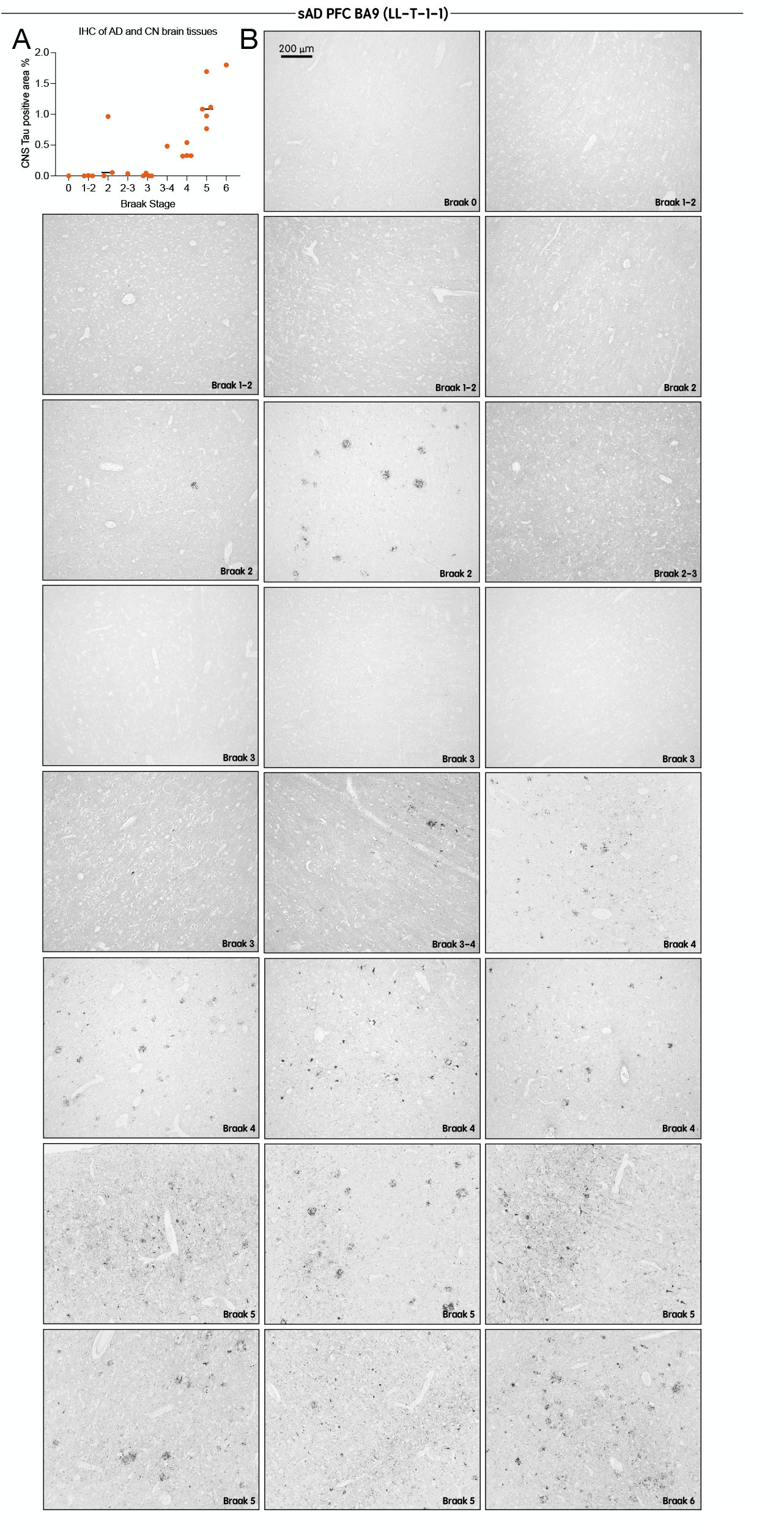
LL-T-1-1 labeling of tau pathology in AD brains reflects Braak stage progression. (A) Quantification of LL-T-1-1 immunohistochemical staining densities in the BA9 region across all Alzheimer’s disease (AD) cases (n = 23). (B) Low-magnification images of LL-T-1-1 immunohistochemistry in the BA9 region of AD brain sections grouped by Braak stages 0–6. LL-T-1-1–positive tau pathology increases with advancing Braak stage, consistent with the characteristic spatial progression of tau pathology in AD.

### CNS-Tau levels in plasma strongly correlate with cognition

To extend CNS-tau quantification to human plasma, we employed NanoMosaic’s label-free, ultra-sensitive nanoneedle detection platform, which utilizes an array of approximately 20,000 nanoneedles (soft-matter material developed by NanoMosaic) to enable high-precision, high-sensitivity quantification of individual biomarkers. The system is implemented in a consumable plate format compatible with Society for Biomolecular Screening (SBS) standards (96-, 384-, or 1536-well formats), facilitating seamless integration with both automated liquid handling systems and manual workflows (Figure. 4A). Nanoneedles are densely patterned on the bottom surface of each well and imaged simultaneously using a complementary metal-oxide-semiconductor (CMOS) color camera in dark-field microscopy mode, enabling rapid and label-free detection without the need for lasers or high-intensity light sources. A representative high-magnification pre-assay image of the nanoneedle array is shown in Figure. 4B, illustrating a 35 × 35 nanoneedle field. To enable tau detection, capture antibodies were immobilized across the nanoneedle surface. Following sample incubation, a biotinylated detection antibody was introduced to form a sandwich immunoassay format. Signal amplification was achieved using a proprietary mass amplifier reagent that binds to the biotin tag, enhancing the mass loading on nanoneedles (Figure. 4C). A post-assay image was captured and compared with the corresponding pre-assay image to assess colorimetric shifts induced by mass accumulation (Figure. 4D). Signal intensities from ∼20,000 nanoneedles were quantified and converted to concentration values based on a standard curve generated using recombinant full-length Tau-441 protein (Figure. S2). Among the LL-T-1 series antibodies, LL-T-1-5 (instead of LL-T-1-1 due to high background of the latter, Figure. S1A) was identified as the optimal capture antibody when paired with Tau-12 as the detection antibody, based on signal-to-noise ratios across serial dilutions (Figure. S1A). This combination enabled a CNS-tau assay with both the lower limit of quantification (LLoQ) and the limit of detection (LoD) below 1 pg/mL (Fig. S1B). LoD was defined as the lowest concentration yielding a signal >2.5 standard deviations above the blank, and LLoQ as the lowest concentration with a coefficient of variation (CV) <20% and recovery within ±20%. Analytical performance of the assay was robust, with an average CV of 8.59% across the standard curve (0.39–100 pg/mL). Excellent dilution linearity was observed in plasma samples (2×, 4×, 8× dilutions into assay buffer), with r^2^ values of 0.93 and 0.99 in representative healthy control and Alzheimer’s disease plasma samples, respectively (n = 1 each for validation) (Fig. S2C). In the current study, we analyzed 72 plasma samples from participants in our BWH memory clinic cohort. Plasma samples were assayed in triplicate at a 3× dilution, yielding an average CV of 7.7%, consistent with the performance observed in standards.

**Figure 4.**
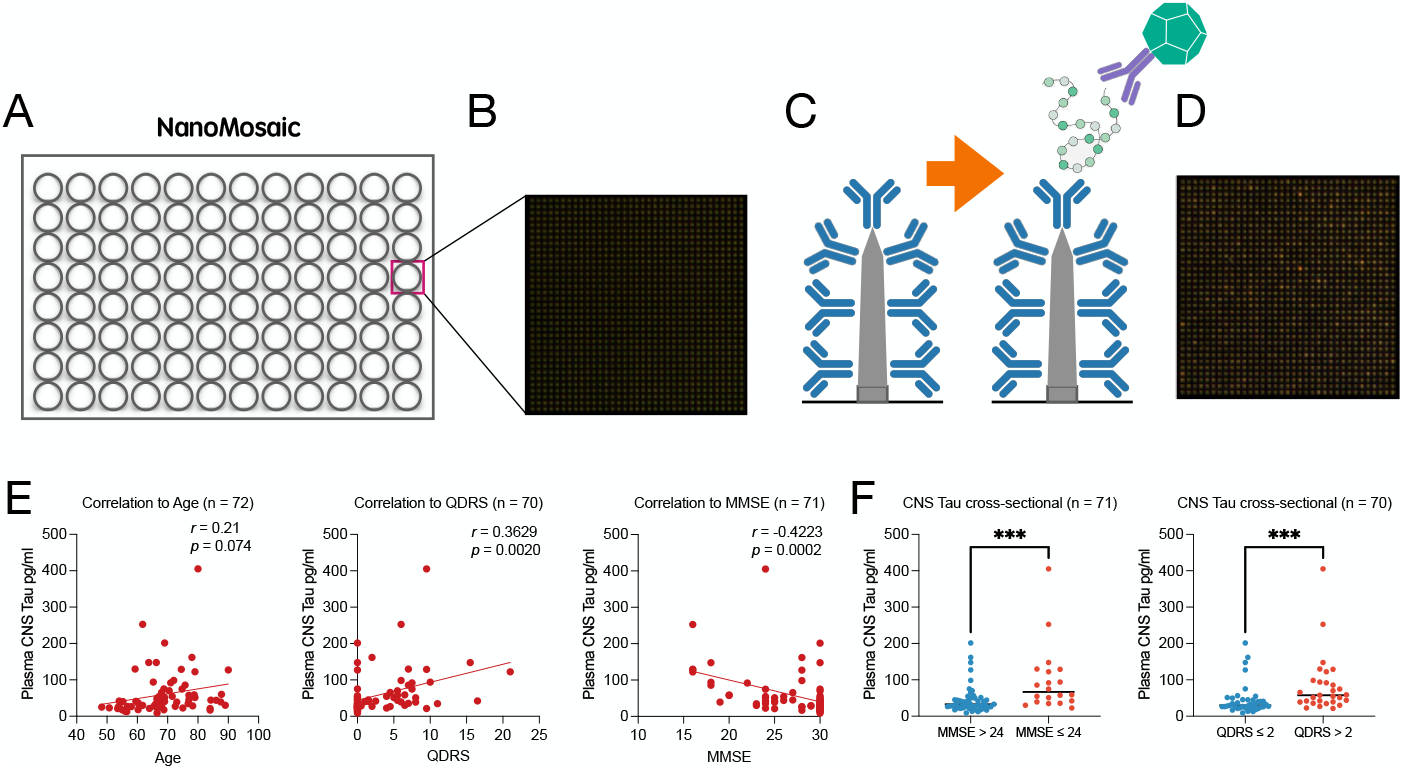
Plasma CNS-tau levels correlate with cognitive impairment. (A) Schematic of the nanoneedle array integrated into a 96-well plate format. (B) High-magnification dark-field image of the nanoneedle surface prior to assay. (C) Illustration of the sandwich immunoassay, with LL-T-1-5 antibody immobilized on each nanoneedle for target capture, followed by detection using labeled secondary antibodies. (D) Representative post-assay image showing colorimetric signal changes on the nanoneedle array compared to (B). (E) Correlation of plasma CNS-tau levels with (left to right) age, QDRS score, and MMSE score. Pearson’s correlation coefficients were used for statistical analysis. (F) Cross-sectional comparisons of plasma CNS-tau levels between cognitively impaired and unimpaired individuals, classified by MMSE (< 24) or QDRS (> 2). Group differences were assessed using the non-parametric Mann–Whitney U test. ***p < 0.001.

We found that the plasma CNS-tau levels in this cohort did not correlate with age, in contrast to other tau species such as NT1-tau (ref). However, CNS-tau levels were significantly correlated with cognitive state as measured by both the Quick Dementia Rating System (QDRS) and the Mini-Mental State Examination (MMSE) (Fig. 4E). Furthermore, when participants were stratified based on MMSE (<24) or QDRS (>2) thresholds, plasma CNS-tau levels were significantly elevated in cognitively impaired individuals (Fig. 4F).

## Discussion

The distinction between CNS-tau and PNS-tau splicing isoforms has been debated for over three decades^6,9^, yet functional and pathological comparisons have been limited^10^. With the recent emergence of blood-based tau biomarkers for AD, the field has revisited this foundational question. Key knowledge gaps persist: (1) the anatomical distribution and relative abundance of CNS- vs. PNS-tau isoforms in human tissues; (2) whether PNS-tau contributes to tauopathy pathogenesis^11^; and (3) how these isoforms are differentially secreted or released into CSF and plasma. Addressing these questions requires precise analytical tools capable of distinguishing isoform-specific signals in both experimental systems and clinical specimens.

In this study, we report the first fully validated CNS-tau specific recombinant monoclonal antibodies, the LL-T-1 series, which have nanomolar affinity and exceptional isoform specificity. The use of recombinant monoclonal antibodies offers a major advantage by eliminating batch-to-batch variability, ensuring consistent performance across experiments and over time, which is critical for both analytical validation and clinical translation. Unlike existing commercial antibodies that broadly recognize multiple tau isoforms or show cross-reactivity, the LL-T-1 series consistently demonstrated selective binding to CNS-tau across diverse platforms: biolayer interferometry, immunoblotting, ELISAs, and immunohistochemistry. This specificity enabled us to differentiate CNS-tau expression from PNS-tau in cultured cells and to visualize selective tau pathology in human brain sections.

Importantly, LL-T-1-1 revealed a distinct pathological pattern in sporadic AD brains. It preferentially labeled dystrophic neurites and neuronal threads over mature neurofibrillary tangles, a profile that differs from widely used phospho-tau antibodies that recognize both CNS-tau and PNS-tau such as AT8 or pT217^5^. This may partially explain why CNS-tau levels in plasma, measured using LL-T-1 antibodies in our study with IHC, correlate more closely with amyloid-related neurodegeneration than with classical tangle pathology as demonstrated by previous report^3^. In contrast, in primary tauopathies such as progressive supranuclear palsy (PSP) and corticobasal degeneration (CBD), LL-T-1-1 robustly stained all hallmark tau inclusions— including tufted astrocytes, coiled bodies, and astrocytic plaques—highlighting its broader utility in tauopathy diagnosis beyond AD.

The translational potential of the LL-T-1 series was further supported by its performance in fluid biomarker assays. Plasma CNS-tau levels measured via an ultrasensitive label-free NanoMosaic platform correlated significantly with clinical cognitive measures (MMSE and QDRS) in real-world memory clinic patients. These findings reinforce the relevance of CNS-tau as a blood-accessible marker of neuronal dysfunction and suggest that its measurement could complement or refine current p-tau–based plasma biomarkers.

This study also offers a novel framework for dissecting tau isoform biology *in vivo*. By leveraging isoform-specific antibodies, future research may address outstanding questions surrounding dynamic tau mechanisms, including whether CNS- and PNS-tau differ in their susceptibility to aggregation, proteolysis, or clearance. Notably, the differential immunoreactivity of LL-T-1-1 across Braak stages implies that CNS-tau–positive pathology evolves with disease progression—consistent with stage-specific vulnerabilities in the alteration of cortical tau. Looking forward, we aim to expand our antibody toolkit by generating PNS-tau– specific antibodies, targeting the exon 4a–encoded region unique to PNS-tau. This will enable direct comparisons of CNS- vs. PNS-tau expression, trafficking, and biomarker relevance. We also plan to assess the clinical utility of CNS-tau assays in PSP and CBD cohorts, where fluid biomarkers remain underdeveloped despite a clear need for improved diagnostics and therapeutic monitoring^12^.

In summary, our findings establish the LL-T-1 series as a novel CNS-tau–specific antibodies with broad applications for neuropathological characterization and fluid biomarker development. The ability to distinguish CNS-tau from PNS-tau opens new avenues for mechanistic studies of tau pathology and holds promise for more refined, disease-specific diagnostic tools across the spectrum of tauopathies.

## Supporting information

Figur S1

## Acknowledgments

Brain tissue samples included in these studies were obtained from Department of Pathology of Brigham and Women’s Hospital and the Harvard Brain and Tissue Resource Center, an NIH NeuroBioBank site. The authors express gratitude to all brain donors and their families for their generosity.

## Declaration of Interests

Luke Slominski and Qimin Quan are employees of NanoMosaic, Inc. DJS is a director of Prothena Biosciences and an ad hoc consultant to Eisai and Roche. All other authors have nothing to disclose.

**<u>Supplementary Figure 1. Development of the plasma CNS-tau assay using the NanoMosaic platform</u>**. (A) Comparison of three LL-T-1 clones used as capture antibodies (paired with Tau-12 as detector) for recombinant Tau-441, evaluating signal-to-noise ratios. (B) Standard curve generated using LL-T-1-5 as the capture and biotinylated Tau-12 as the detector antibody, calibrated with recombinant Tau-441. (C) Dilution linearity of the plasma CNS-tau assay across two human plasma samples at 2×, 4×, and 8× dilutions, demonstrating strong linear response.

