## Supplementary figures and images for "CNS-Tau Specific Antibodies Illuminate Disease Signatures Across Tauopathies"

### Figur S1

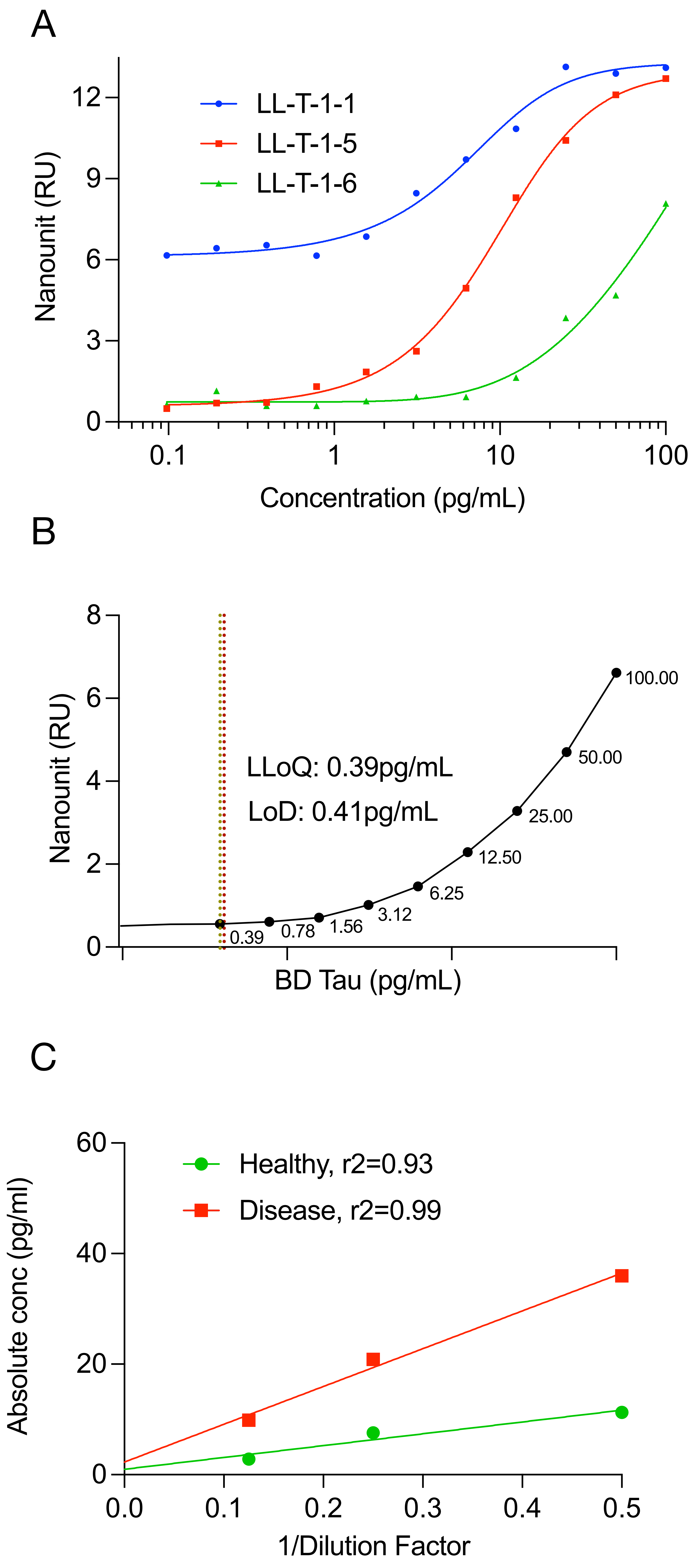
